# Synthetic methanogenic communities reveal differential impact of ecological perturbations on aceto- and hydrogeno-trophic methanogens

**DOI:** 10.1101/307041

**Authors:** Jing Chen, Matthew Wade, Jan Dolfing, Orkun S Soyer

## Abstract

Synthetic microbial communities provide reduced microbial ecologies that can be studied under defined conditions. Here, we use this approach to study the interactions underpinning anaerobic digestion communities and involving the key microbial populations of a sulfate reducer (*Desulfovibrio vulgaris*), and aceto-(*Methanosarcina barkeri*) and hydrogenotrophic (*Methanococcus maripaludis*) methanogens. We create all possible mixed culture combinations of these species and analyse the stability and productivity of each system over multiple sub-culturings and under different sulfate levels, mimicking ecological perturbation in the form of strong electron acceptor availability. We find that all three species can co-exist in the absence of sulfate, and that system productivity (in form of methane production from lactate) increases by almost two-fold compared to co-cultures. With increasing sulfate availability, co-existence is perturbed and both methanogenic populations display a diminishing trend. Interestingly, we find that, despite the continued presence of acetate in the system, the acetotrophic methanogens are more readily disrupted by sulfate perturbation. We show that this is due to a shift in *M. barkeri* metabolism towards increased co-utilisation of hydrogen with acetate, which we verified through experiments on mono cultures and mass balance calculations in co-cultures. We conclude that hydrogen is a key factor for both hydrogeno- and aceto-trophic methanogenesis and can influence these populations differentially under the common ecological perturbation of strong electron acceptor availability. These findings will help engineering of larger synthetic communities for specific applications in biodegradation and understanding complex anaerobic digestion communities found in animal guts, sediments, and bioreactors.

## INTRODUCTION

Microbial communities constitute the dominant mode of living for microorganisms in nature. All studied habitats, ranging from human and animal guts, to the soil and ocean, are found to be inhabited by microbial communities composed of hundreds of different species (Widder *et al.*, 2016). Interactions among these species ultimately give rise to community-level functions, including metabolic conversions that enable animal and plant nutrition (Bulgarelli *et al.*, 2013; Lee and Hase, 2014), and geo-biochemical cycles (Falkowski *et al.*, 2008; Fuhrman *et al.*, 2015). Understanding the biochemical and physical basis, and the ecological and evolutionary drivers of functional stability in microbial communities is thus a key open challenge in microbial ecology (Widder *et al.*, 2016). Achieving better understanding of these drivers for stable community function can enable prediction of functional stability and collapse thereof (Allison and Martiny, 2008; Becker *et al.*, 2012), the design of interference strategies to shift community function (Shetty *et al.*, 2017; Zerfaß *et al.*, 2018), and the engineering of bespoke ‘synthetic communities’(Haruta *et al.*, 2013; De Roy *et al.*, 2014; Grosskopf and Soyer, 2014; Lindemann *et al.*, 2016).

Towards deciphering ecological and evolutionary drivers of function and functional stability in microbial communities, methanogenic anaerobic digestion (AD) offers an ideal model system, where production of methane from complex organic substrates can be taken as a proxy for community function. AD communities are found in many environments including ocean and lake sediments, soil, and animal guts, and are utilised in biotechnological revaluation of organic waste (Spirito *et al.*, 2014). It is well known that high substrate levels and limited availability of electron acceptors in the AD system can create thermodynamic limitations that can dominate functional stability and community dynamics (De Vrieze and Verstraete, 2016), underpin the emergence and maintenance of diversity in the community (Grosskopf and Soyer, 2016), and drive evolution of metabolic interactions among different species (Schink, 1997; Embree *et al.*, 2015). A key reason for the importance of thermodynamic limitations in AD systems is that it forces a cooperative (i.e. syntrophic) metabolism of organic acids, whereby degradation of these compounds by one group of organisms can only be maintained (i.e. be thermodynamically feasible) by continuous removal of end-products by another (Schink, 1997; Stams and Plugge, 2009). This syntrophic degradation can be performed by a range of fermentative microbes including sulfate reducers, while the second step of end-product removal can only be performed by aceto- and hydrogeno-trophic methanogens, which specialise in the consumption of acetate and hydrogen, respectively (Schink, 1997; Thauer *et al.*, 2008). In the case where the syntrophic degradation step is disrupted, acetate and hydrogen can accumulate, leading to further thermodynamic inhibition, as well as acidification, ultimately causing the functional collapse of the AD system (Demirel and Scherer, 2008; Wang *et al.*, 2018).

Despite the importance of syntrophic interactions between methanogens and secondary degraders, our understanding of ecological and evolutionary factors influencing these interactions is still limited. Among the ecological factors, it is known that syntrophy can be destabilized by an increased availability of strong electron acceptors, such as nitrate and sulfate. These electron acceptors not only shift metabolism of secondary degraders towards respiration, but also allow them to utilise acetate and hydrogen (Badziong *et al.*, 1979; Noguera *et al.*, 1998), potentially causing competitive exclusion of methanogens that rely solely on these substrates (Thauer *et al.*, 2008; Paulo *et al.*, 2015). It is shown that changes in carbon dioxide and hydrogen partial pressures can also influence syntrophic interactions among acetate oxidising bacteria, and aceto- and hydrogeno-trophic methanogens, primarily through changes in the energetics of key metabolic reactions (Mayumi *et al.*, 2013; Kato *et al.*, 2014). Many studies, however, show that both aceto- and hydrogeno-trophic methanogenesis can still co-exist with secondary degraders in the presence of significant concentrations of strong electron acceptors (Whiticar *et al.*, 1986; Kuivila *et al.*, 1990; Dar *et al.*, 2008), and can persist or adapt to perturbations in form of electron acceptor addition (Raskin *et al.*, 1996; Ma *et al.*, 2017).

Besides the general view that strong electron acceptor availability can create a competition between methanogens and other species, the specific effect of this ecological perturbation on the different methanogenic groups remains an open question (Ozuolmez *et al.*, 2015). In particular, methanogens are distinguished into two major groups through their respiratory and energy-conserving mechanisms (Thauer *et al.*, 2008; Kulkarni *et al.*, 2009; Ferry, 2010), which has a bearing on their ability and preference to utilise H_2_ and acetate (Thauer *et al.*, 2008). Acetotrophic methanogens belong to the group that encodes key respiratory cytochromes that allow them to utilise acetate (and for some species also other methyl-containing single carbon molecules) (Thauer *et al.*, 2008; Kulkarni *et al.*, 2009). It has been shown that some of these cytochrome encoding, acetotrophic methanogens maintain the ability for hydrogenotrophic methanogenesis, and can also co-utilise H_2_/CO_2_ with other single carbon molecules including acetate (Muller *et al.*, 1986; Thauer *et al.*, 2008; Kulkarni *et al.*, 2009). It is unclear if and how this flexibility in methanogenic pathways allows acetotrophic methanogens to withstand ecological perturbations, compared to obligate hydrogenotrophic methanogens.

Here, we study this question using a synthetic community construction approach and focusing on specific sulfate reducing bacteria, and aceto- and hydrogeno-trophic methanogens. The competition between these different functional groups under sulfate availability has not been studied in a defined community before, prompting us to create and analyse mono-, co-, and tri-cultures of representative species from these functional groups. We evaluated productivity and stability in the resulting communities under perturbations in the form of sulfate availability, representing a strong electron acceptor. This revealed that, in the absence of sulfate, inclusion of both of the methanogenic populations increases methane production from lactate by almost two-fold compared to co-cultures of the sulfate reducer with a single methanogen. With increasing sulfate availability, however, we find a differential impact on the two methanogenic groups. While hydrogenotrophic methanogens were lost from the community at sulfate levels that only allow full respiration of the available lactate, acetotrophic methanogens were lost readily at lower sulfate levels. This differential stability was also evident at the level of productivity in the tri-culture, where contribution from acetotrophic methanogenesis reduced with increasing sulfate. These results on stability and productivity could be explained through mass balance calculations, but only if we assumed a dependency of the acetotrophic methanogen on hydrogen. We have then verified this assumption experimentally using monocultures. Together, these results show that hydrogen-based competition in presence of strong electron acceptors can influence both aceto- and hydrogeno-trophic methanogens, with the former being more prone to be lost from the system as a result. These findings are of significant relevance to understand complex, natural AD communities, and to further engineer synthetic communities mimicking their functionality and optimised for specific applications.

## RESULTS

To better understand the functional role and stability of syntrophic interactions between sulfate-reducing bacteria and methanogens in AD communities, we created here a set of synthetic microbial communities composed of two and three species. We used three key species to represent the roles of sulfate-reducing bacteria (*Desulfovibrio vulgaris; Dv*), and aceto- (*Methanosarcina barkeri; Mb*) and hydrogeno-trophic methanogens (*Methanococcus maripaludis; Mm*). The *Dv-Mm* pair has emerged in recent years as a model system to study syntrophic interactions (Hillesland and Stahl, 2010), while *Mb* is one of the most well-studied methanogens capable of acetotrophic methanogenesis (De Vrieze *et al.*, 2012). We cultivated these organisms using a common, defined media and created each possible synthetic community composed of one, two, and three species (see *Methods*). We initiated replicate synthetic communities using a chemically-defined media with lactate (30 mM), as the sole organic carbon source, and cultivated them under different levels of sulfate (see *Methods*). Each constructed community was incubated, and sub-cultured twice, over three-week periods. These conditions mimicked a low-flow, chemostat-like system, while different levels of sulfate mimicked different availability of strong electron acceptors.

### All species co-exist and community productivity increases in the absence of strong electron acceptors

Presence of both methanogenesis routes through aceto- and hydrogenotrophic species is expected to increase productivity in AD communities due to a more complete conversion of the key fermentation products from secondary degradation (Fig. 1A). We found this expectation to be fulfilled in the absence of sulfate; the synthetic *Dv-Mm-Mb* tri-culture produced close to 2-fold more methane compared with the *Dv-Mm* and *Dv-Mb* co-cultures (Fig. 1B). The tri-culture and the *Dv-Mm* co-culture achieved stable methane levels over three sub-cultures, while methane production in the *Dv-Mb* co-culture was highly variable. In line with these observations, the tri-culture and the *Dv-Mm* system displayed full lactate conversion, while there was significant lactate remaining in one replicate *Dv-Mb* system (Fig. 1C). Interestingly, both the tri-culture and the *Dv-Mb* co-culture displayed also significant levels of residual acetate, indicating that *Mb* is not able to consume all of the acetate fermented by *Dv* (Fig. 1D). This finding was replicated when we cultivated the cultures under a five-week sub-culturing regime (Fig. S1), suggesting that lack of full acetate consumption is not simply due to slow growth of *Mb* on this substrate.

**Figure 1.**
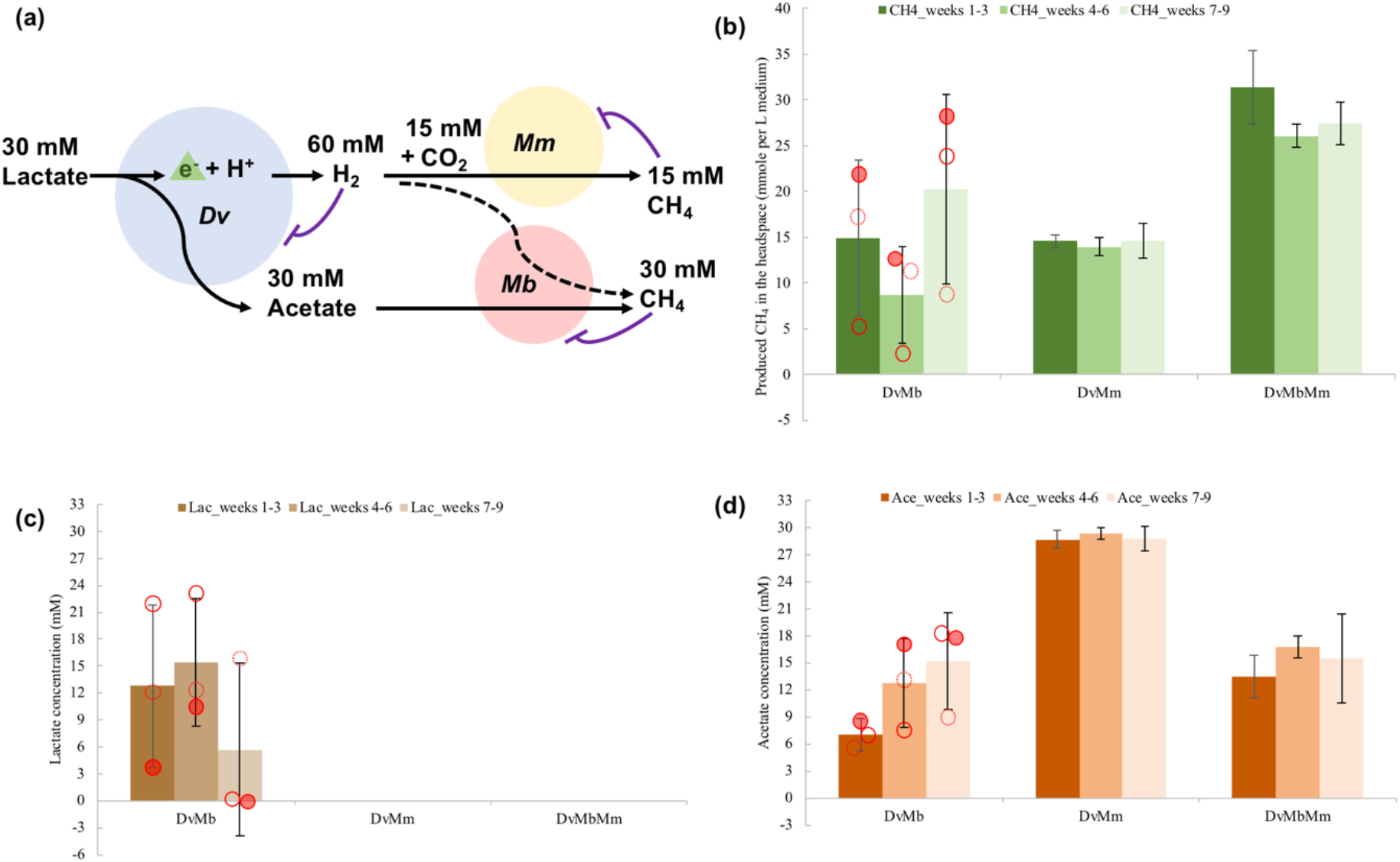
**(a)** Schematic of possible interactions of the three species for converting lactate to methane. The three different species *Desulfovibrio vulgaris* (*Dv*), *Methanococcus maripaludis* (*Mm*) and *Methanosarcena barkeri* (*Mb*) are shown as blue, yellow, and red circles respectively. The metabolite concentrations shown are those based on the stoichiometries of reactions given in Table 2 and using 30mM initial lactate. Possible thermodynamic inhibitions are indicated by t-ended arrows. The dashed line indicates possible co-utilization of H_2_ by *Mb*. **(b)** Methane produced in the headspace in the absence of sulfate and in the different co- and tri-cultures as indicated on the x-axis. Measurements from 5 mL test tube cultures are used to extrapolate to 1 L culture output, so to achieve a better comparison of gas and organic acid data (in mM). **(c, d)** Lactate and acetate detected in the liquid phase after 21 days cultivation without sulfate addition. Red dots in these two panels refer to the three replicates in the *Dv-Mb* co-cultures. (replicate 1 – red hollow circle, replicate 2 – dashed red hollow circle, and replicate 3 – filled red circle). Error bars on panels b-d are based on three replicates.

### Increased electron acceptor availability shows differential impact on the maintenance and productivity of aceto- and hydrogeno-trophic methanogens

In order to find out the impact of sulfate availability on the stable co-existence of sulfate-reducing bacteria and different methanogens, we further analysed the dynamics of each co-culture and the triculture at different sulfate levels. In particular, we cultivated communities in sulfate concentrations that provide either half or full stoichiometric equivalence to lactate; i.e. 7.5 or 15mM sulfate allowing either half or full respiration of lactate by *Dv* (these conditions are referred to as ‘half-’ and ‘full-sulfate’ from now on). We found that increased sulfate availability immediately impacted the *Dv-Mb* co-culture and resulted in a loss of methane production already in half-sulfate treatments (Fig. 2). The *Dv-Mm* co-culture displayed stable co-existence at half-sulfate treatments, but methanogenesis was clearly showing a diminishing trend in the full-sulfate treatment (Fig 2). Methanogenesis under increasing sulfate levels in the synthetic tri-culture behaved qualitatively similarly to the *Dv-Mm* coculture, but methane levels in the tri-culture during each culturing period were slightly higher (Fig. 2).

**Figure 2.**
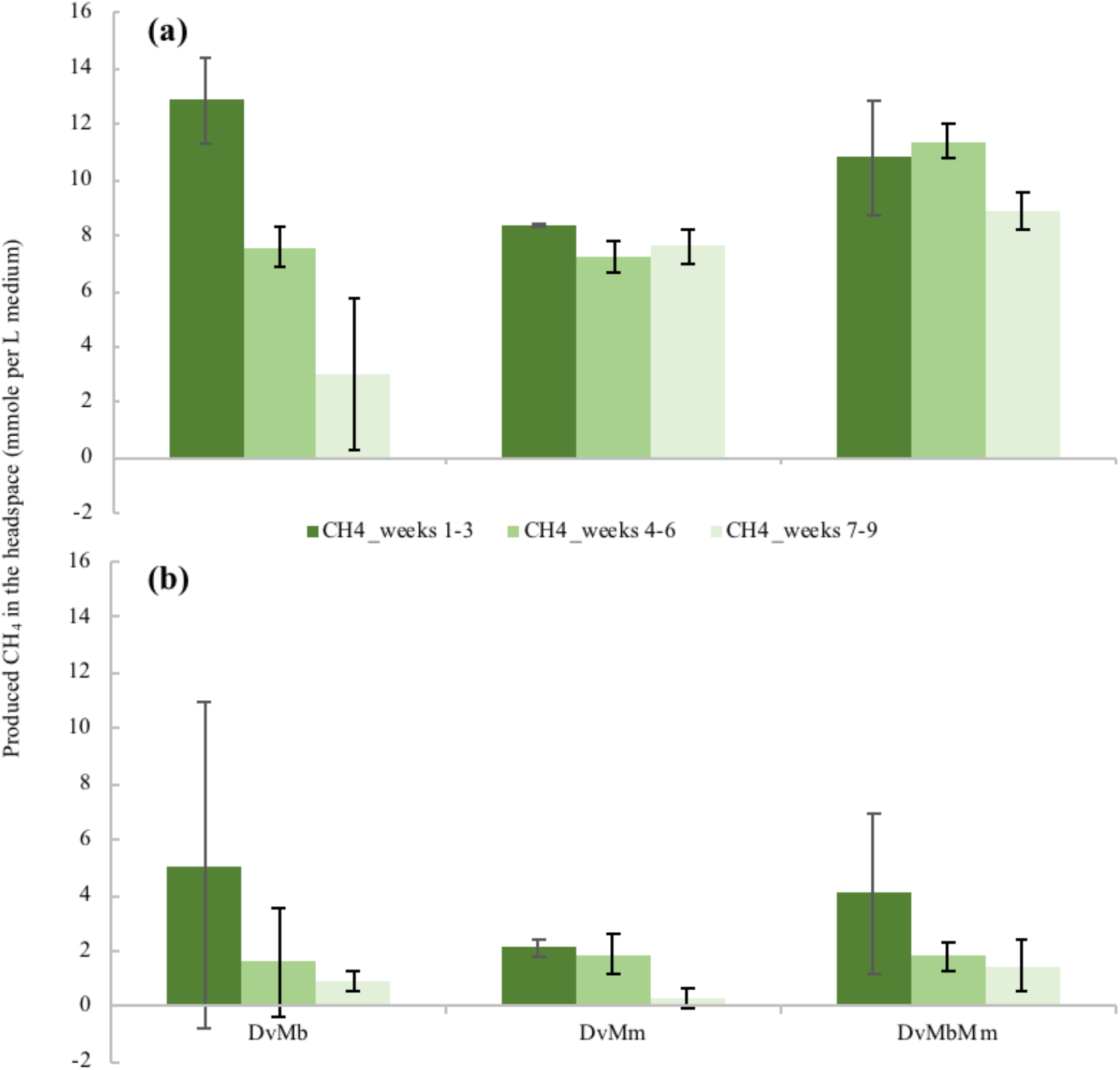
Produced methane in the headspace after 21 days cultivation with 7.5 mM **(a)** and 15 mM **(b)** sulfate addition. The different co- and tri-cultures composed of species *Desulfovibrio vulgaris* (*Dv*), *Methanococcus maripaludis* (*Mm*) and *Methanosarcena barkeri* (*Mb*), as shown on the x-axis. Measurements from 5 mL test tube cultures are used to extrapolate to 1 L culture output, so to achieve a better comparison of gas and organic acid data (in mM; see also Fig. S2). Colors indicate different culturing periods as shown in the legend. Error bars on both panels are based on three replicates.

We found that the impact on methane production by switching from individual cocultures to a tri-culture, also depends on sulfate availability (compare Fig. 1B and Fig. 2). In the absence of sulfate, the tri-culture produced close to 100% more methane compared with the co-cultures, instead, the difference in the methane production between these two groups was 32.28% under the half-sulfate treatment. This suggests that *Mb* populations are either diminishing under the half-sulfate treatment or are not receiving enough acetate. We excluded the latter possibility by measuring lactate and acetate levels for all co-cultures and the tri-culture, and under each sulfate treatments (Fig. S2). This showed that there are significant levels of acetate in the tri-culture under half-sulfate treatment (as well as full-sulfate treatment), suggesting that the observed smaller increase in productivity (from co- to tri-cultures) compared to the no-sulfate case is not due to acetate limitation.

To further corroborate these findings, we analysed community stability at the species level by enumerating the different populations using quantitative PCR (qPCR) of the targeted species gene copies at the end of the overall experiment (see *Methods*). In general, *Dv* populations accounted for a large fraction (>80%) of the overall community in all treatments and displayed an increasing trend with sulfate addition (Fig. S3A). An opposite trend is observed for the population sizes of *Mm* and *Mb*, as expected from the observed decrease in methane production. The *Mb* abundance showed high variance in most cases, except for the tri-culture with no sulfate, while *Mm* populations showed an increase in tri-culture (for all distinct sulfate treatments) compared to the corresponding co-culture (Fig. S3B). Taken together, these findings suggest an increased stability of methanogen populations with the increased community complexity (i.e. extended syntrophic interactions) both under sulfate perturbation and without sulfate, and a lower stability of *Mb* populations compared to *Mm*, as sulfate becomes available.

### *Mb* populations productivity from acetate shows significant dependence on H_2_

Why can the acetotrophic *Mb* contribute to methane production under no-sulfate treatment, but not under half- and full-sulfate treatments, even though there is enough acetate available for it to grow? As shown above, *Dv* contributes to a higher fraction of the population with increasing sulfate, and can utilize H_2_, as well as lactate, under this condition (Noguera *et al.*, 1998). This creates a competitive situation for *Mm*, but possibly also for *Mb*, if it relies also on H_2_ for maintaining its population size. Indeed, we observed H_2_ utilization by *Mb* both in control mono-cultures, with lactate as the sole carbon source (Fig. S4), as well as in two replicates in the final sub-culturing of the *Dv-Mb* co-cultures under no-sulfate treatment (Fig. S5).

These observations, as well as previous indications of H_2_ utilisation of *Mb* (Muller *et al.*, 1986; Thauer *et al.*, 2008; Kulkarni *et al.*, 2009), prompted us to more directly test the impact of H_2_ on the growth of *Mb* with acetate, using its monocultures (see *Methods*). These experiments showed that, with acetate provided at 30mM, increasing H_2_ pressure in the headspace significantly increased *Mb*’s methane production (Fig. 3). Although most acetate was consumed both in the presence and absence of H_2_, the methane production under the latter condition was only the third of that in the presence of 80% H_2_ in the headspace; 20 mM vs. 60 mM methane, respectively. The 1:2 stoichiometric relation between acetate and methane in presence of 80% H_2_ in the headspace, suggests that under this condition, *Mb* utilizes H_2_ oxidation with acetate reduction, as well as, or in place of, acetotrophic methanogenesis.

**Figure 3.**
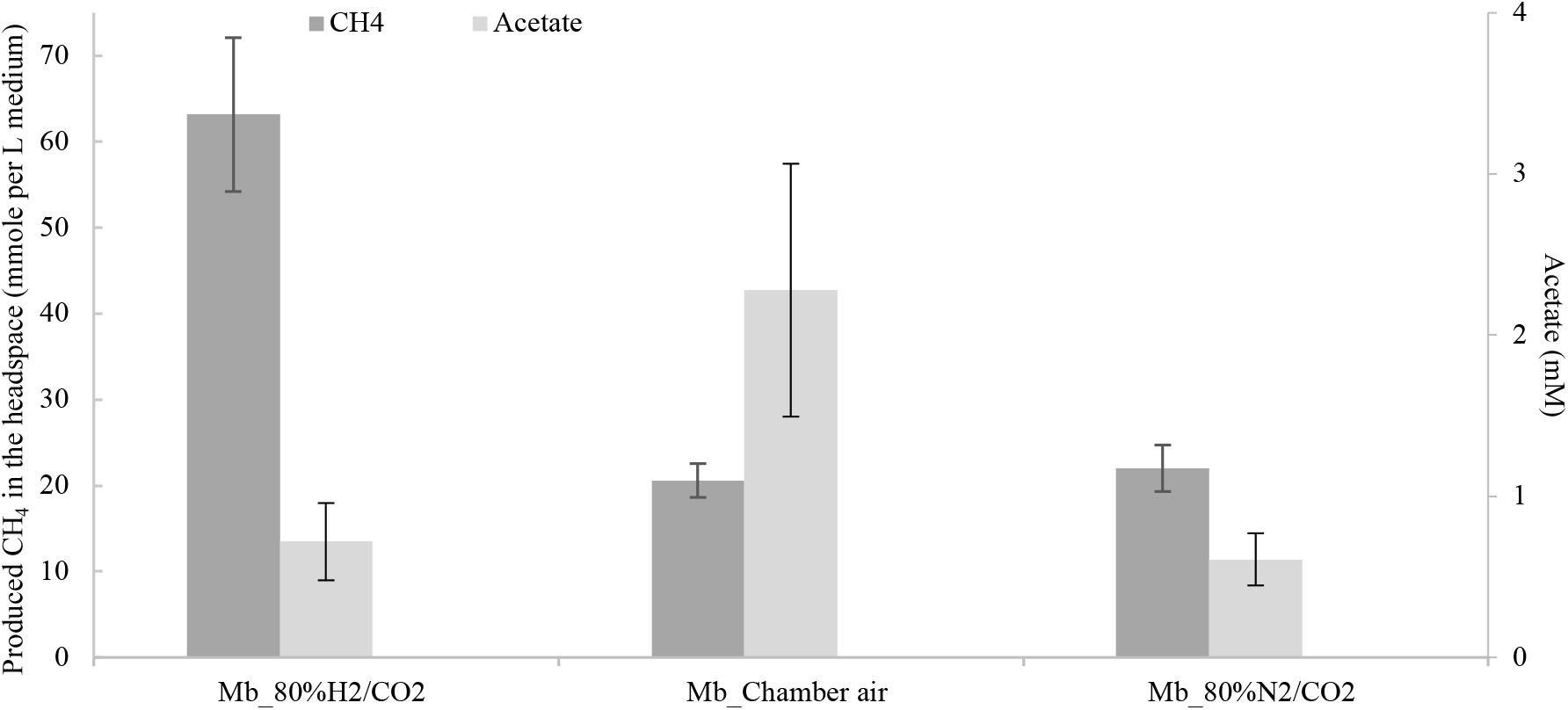
Produced methane in the headspace by *Methcmosarcinci barkeri* (*Mb*) monocultures (dark gray) and detected residual acetate (light gray) after three weeks of cultivation with 30mM initial acetate concentration. Note that methane and acetate levels are shown on different y-axes. The three sets of results indicate the initial gas composition in the headspace, as shown on the x-axis; 80% H_2_ (with 20% CO_2_), 3.14% H_2_ (anaerobic chamber, remaining atmosphere is approx. 89.92% N_2_ and 5.32% CO_2_), and 0% H_2_ (with 80% N_2_ and 20% CO_2_ atmosphere) Error bars are based on three replicates.

### Mass balance calculations confirm *Mb*’s use of H_2_ in *Dv-Mb* co-cultures

To further evaluate this observation of H_2_ (co)utilisation by *Mb* mono-cultures in the context of the synthetic communities, we performed mass balance calculations using experimental data from the *Dv-Mb* co-cultures without sulfate addition (Table 1) and the key reactions possible in the system (Table 2). Using initial (30mM) and residual lactate concentrations observed at the end of a three-week cultivation, we derived the observed change in lactate (ΔLactate_obs_). We used this value to calculate the theoretical stoichiometric H_2_ and acetate output by *Dv*, assuming full fermentation of lactate by *Dv* (reaction 5 in Table 2). We combined these calculated levels with the observed ones (change in headspace H_2_ and residual acetate) to then estimate the theoretical H_2_ and acetate levels that would have been available for *Mb* consumption (H_2*Mb*_ and Acetate_*Mb*_; see Table 1). For example, in one replicate (row 1 in Table 1), we found 20.07mM residual lactate, indicating 9.93mM of lactate consumed by *Dv*, resulting in the estimation of acetate and H_2_ production at 9.93 and 19.86mM respectively. For this same example replicate, the observed residual acetate was 6.94mM and headspace H_2_ increased by 2.72mM from its original level, resulting in the estimation of *H*_2*Mb*_ and Acetate_*Mb*_ at 17.14 and 2.99mM.

**Table 1.**
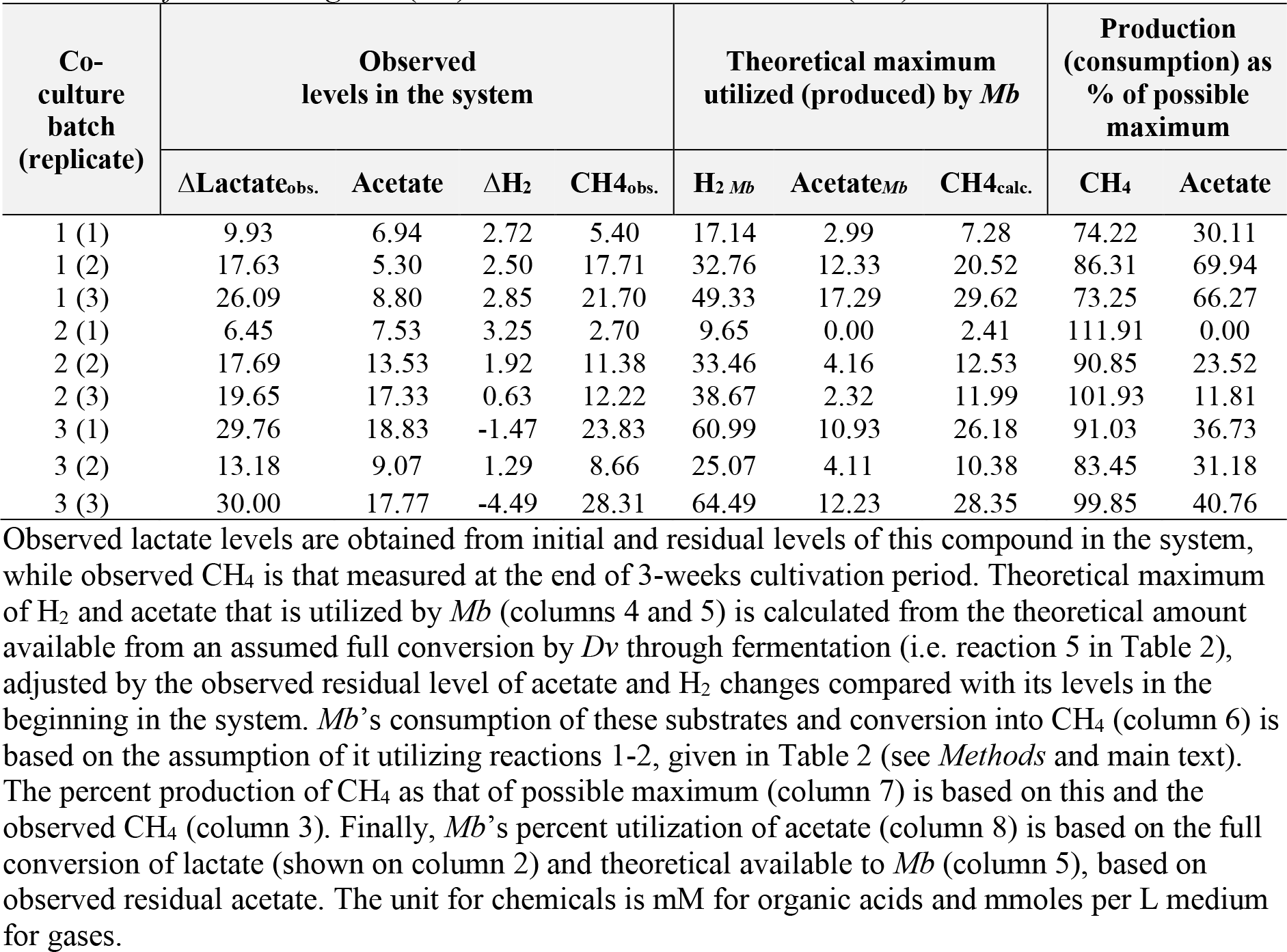
Observed and calculated substrate levels and percent consumption and productions in the *Desulfovibrio vulgaris* (*Dv*) - *Methanosarcina barkeri* (*Mb*) co-culture without sulfate.

**Table 2.**
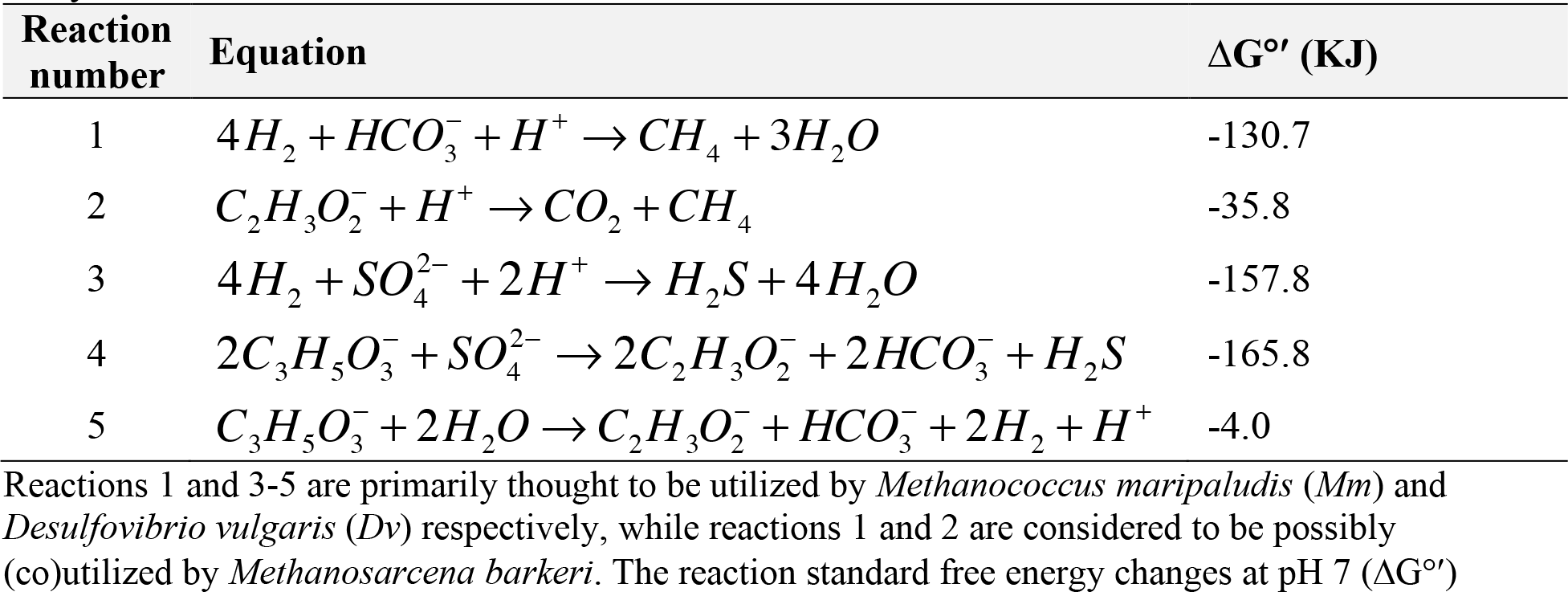
The compounded, overall growth-supporting reactions considered in the present study.

The consumption of these substrates by *Mb* can proceed theoretically through aceto- and hydrogeno-trophic methanogenesis (reactions 1 and 2 in Table 2), and their possible combination through H_2_ oxidation with acetate reduction. If we use H_2*Mb*_ and Acetate_*Mb*_ as given constraints, we can show that the theoretical overall methane output (CH_4calc._) would always be equal to H_2*Mb*_/4 + Acetate_*Mb*_ (see *Methods*). We find that the observed methane in the system (CH_4obs._) was almost always below this theoretical maximum (see Table 1). There were, however, two cases that result in more methane than theoretically possible, by 1% and 10% more. We find that these two cases present the lowest acetate consumption (no detectable consumption in the second case), and the highest H_2_ consumption, indicating significant H_2_ consumption by *Mb* to produce methane through reaction 1 (and possibly also combination of reactions 1 and 2). This might have altered *Dv*’s metabolism to shift from acetate fermentation into H_2_ production (Walker *et al.*, 2009; Grosskopf *et al.*, 2016) and/or its investment of reductive power into biomass production, which could explain the discrepancy with our theoretical calculation based on reaction 5.

Overall, our results summarised in Table 1 show that the methane production in the system cannot be explained solely by acetotrophic methanogenesis but requires involvement from reactions 1 and 2, or their combination. Note that this general conclusion would not be affected by possible investment into biomass by *Dv* or *Mb*, which we neglected in the calculations shown in Table 1. Moreover, methane production as percentage of the theoretical maximum (as calculated above) increases over the course of the three sub-culturing periods, while acetate consumption decreases (Table 1). In other words, *Mb* seems to be shifting its metabolism in the presence of *Dv* in a way favouring increasingly H_2_ (co)utilisation. This trend, in turn, could explain the instability of *Mb* in the co- and tri-cultures under increasing sulfate conditions, where competition for H_2_ would be higher (due to utilisation both by *Dv* and *Mm*).

## DISCUSSION

We have developed for the first time a full set of co- and tri-cultures comprising three key functional populations found in AD systems, a sulfate reducer (*Dv*) and aceto- (*Mb*) and hydrogeno-trophic (*Mm*) methanogens. These systems allowed us to study the syntrophic and cross-feeding interactions among these species under a common ecological perturbation in form of sulfate availability. Our results showed an increased productivity, in the form of methane production, and high stability, through species co-existence, in the tri-cultures with no sulfate addition. With an increasing availability of sulfate, the shift in *Dv* metabolism towards respiration created a disruption in the methanogen populations, and under nonlimiting sulfate concentrations we found both aceto- and hydrogeno-trophic methanogenesis showing a strongly diminishing trend. At limiting levels of sulfate, the disruption to co-existence was also limited, but we found a differentially stronger impact on acetotrophic populations represented by *Mb*. Experiments on the monoculture of this species verified the strong influence of H_2_ on its growth with acetate, suggesting that its observed instability in tri- and co-cultures could be due to competition with *Dv* and *Mm* for this compound.

Perturbation of methanogenic populations due to competition for H_2_ with sulfate reducers has been postulated and studied in several complex communities (Whiticar *et al.*, 1986; Kuivila *et al.*, 1990; Raskin *et al.*, 1996; Dar *et al.*, 2008; Ma *et al.*, 2017). The presented study, with its well-defined, simplified synthetic communities, provides a direct observation of this competition, and instability of methanogens, in the presence of a sulfate reducer and sulfate as an electron acceptor. More importantly, these synthetic communities reveal that acetotrophic methanogens are more prone to suffer from such sulfate-inflicted instability despite their primary substrate being acetate rather than H_2_. It would be very interesting to further evaluate this finding in the context of complex AD communities found in nature and in bioreactors. In particular, there is some evidence from the latter systems that hydrogen supplementation can lead to higher methane production (Bassani *et al.*, 2015) which, according to our findings, could be due to a reduction in the competition for H_2_ and enhanced productivity (and possibly growth) of acetotrophic methanogens.

The synthetic community approach presented here can and should be extended to other species’ combinations. In particular, we note that while *Mb* is capable of growth on acetate, there are several other methanogens in nature that seem to have become obligate growers on this substrate, including those from the genus *Methanosaeta* (Ferry, 2010). It would be very interesting to assess the stability of these obligate acetotrophic methanogens against secondary degraders such as sulfate reducers, and hydrogenotrophic methanogens. To this end, a representative species (*Methanosaeta concilii*) from this functional group has already been studied using a synthetic community approach (Ozuolmez *et al.*, 2015), resulting in the identification of both competitive and cooperative interactions with *Dv* and *Mm*. The biochemical underpinning of these interactions, both in that study and the current one, is the flexibility and efficiency of energy conservation mechanisms found in the methanogens (Thauer *et al.*, 2008). Recent studies have shown that the ability to encode different cytochromes and hydrogenases allows *Mb* (and other methanogens encoding cytochromes) to channel electrons resulting from both the oxidation of one-carbon molecules and H_2_ into the reduction of the key heterodisulphide CoM-S-S-CoB (Ferry, 2010). The resulting electron flow scheme allows *Mb* to perform both aceto-and hydrogeno-trophic methanogenesis with higher ATP yield, but causes a higher H_2_ pressure requirement for the latter process compared to obligate hydrogenotrophic methanogens (Thauer *et al.*, 2008). In addition, acetate and one-carbon consumption under this electron flow scheme is suggested to involve H_2_ cycling, whereby H_2_ is generated in the cytosol to then diffuse out of the cell and be reutilised at membrane-bound hydrogenases (Kulkarni *et al.*, 2009). Both its high H_2_ requirement for hydrogenotrophic methanogenesis, and its possible reliance on H_2_ cycling for acetotrophich methanogenesis, makes *Mb* vulnerable to ecological perturbances as we shown here.

In this biochemical context, it would be very interesting to see if *Mb* can adapt to co-culturing with *Dv* under a no-sulfate regime and become more tolerant to sulfate-based perturbances. We observe some indication of such possibility, where some of the *Dv-Mb* replicates shifted to significant H_2_ consumption and produced high levels of methane under the no-sulfate treatment. Even in the case of half-sulfate treatment, we found high variance in the *Dv-Mb* co-cultures in terms of productivity, indicating better ability of *Mb* to utilise H_2_. It would be interesting to further evaluate this possibility of *Mb*’s adaptation into a hydrogenotrophic (H_2_/CO_2_) or mixotrophic (H_2_/Acetate) metabolism, and whether the newly identified electron bifurcation mechanisms in hydrogenotrophic methanogenesis pathways of *Mm* (Costa *et al.*, 2013) could also be present in *Mb* or other acetotrophic methanogens. The evolutionary adaptations under phases of stable syntrophy can indeed be a key factor for the emergence and stability of microbial interactions. While our combination of *Dv, Mm*, and *Mb* is not a naturally occurring one and these species have not necessarily undergone co-evolution (except throughout these experiments), there is now increasing evidence that the interplay of evolutionary and ecological dynamics is important for the emergence and stability of microbial interactions (Cavaliere *et al.*, 2017). For example, recent community coalescence studies find dominance of entire AD communities over others (Sierocinski *et al.*, 2017), suggesting co-adaptation among community members being a key drive of productivity and stability. Supporting this view, enriched AD communities are shown to display additional metabolic interactions (particularly auxotrophic interactions) on top of syntrophic interactions (Embree *et al.*, 2015). Evolutionary adaptations are also seen in the *Dv-Mm* co-culture used here; both species are found to accumulate beneficial mutations when co-evolved in the absence of sulfate (Hillesland *et al.*, 2014), and *Dv* populations are found to harbor polymorphisms that directly influence the ability to form a syntrophic interaction with *Mm* (Grosskopf *et al.*, 2016). Thus, natural communities might display evolutionary adaptations that render them more resilient to perturbations than our synthetic systems, and might display auxiliary interactions on top of the metabolic syntrophies and cross-feeding interactions that we observed here.

Besides their value as experimental hypothesis-generating tools, synthetic communities are also suggested to have potential as specific biotechnological applications (Widder *et al.*, 2016). To this end, the co- and tri-cultures presented here can be further expanded with additional functional groups of microbes to attain biotechnologically relevant conversions. It has been suggested for example that energy limited systems presenting thermodynamically driven syntrophic interactions, as well as cross-feeding can provide enhanced productivity compared to mono-culture based bioproduction (Cueto-Rojas *et al.*, 2015). Certain chemical conversions and degradations of complex biomaterials, such as cellulose, cannot be achieved by monocultures, and for the evaluation of these compounds a synthetic community approach, as presented here, will be necessary. Therefore, it would be interesting to expand the tri-culture presented here with primary degraders to allow conversion of complex sugars into methane, as already attempted for cellulose (Kato *et al.*, 2005). We advocate the combined use of ecological, evolutionary, and engineering approaches to the development and further engineering of such synthetic communities, to achieve robust new biotechnological applications and more representative model ecosystems.

## MATERIALS AND METHODS

### Strains and media

*Desulfovibrio vulgaris* Hildenborough (DSM644, Dv-WT), *Methanosarcina barkeri* (DSM800, Mb), and *Methanococcus maripaludis* S2 (DSM2067, Mm) were originally ordered from the public strain centre DSMZ (www.dsmz.de). The particular *Desulfovibrio vulgaris* strain (referred to as ‘*Dv*’ in this text) used in the present work is isolated previously in our laboratory and presents two key genetic mutations that allow it to grow syntrophically with *Methanococcus maripaludis* without sulfate (Grosskopf *et al.*, 2016). A defined anaerobic medium, adapted from previous studies (Walker *et al.*, 2009; Grosskopf *et al.*, 2016), is used to grow *Dv, Mm*, and *Mb*. The recipe and preparation protocols of this medium are as follows; Basal salt mix: In 1 L dH_2_O, dissolved: K_2_HPO_4_: 0.19 g, NaCl: 2.17 g, MgCl_2_ × 6H_2_O: 5.5 g, CaCl_2_ × 2H_2_O: 0.14 g, NH_4_Cl: 0.5 g, KCl: 0.335 g, NaHCO_3_: 2.5 g. Trace element solution (100X): In 850 mL of dH2O, dissolved 1.5 g Nitrilotriacetic acid and adjust the pH to 6.5 with KOH. Then added: MgCl_2_ × 6H_2_O: 2.48 g, MnCl_2_ × 4 H_2_O: 0.585 g, NaCl: 1 g, FeCl_2_ × 4 H_2_O: 0.072 g, CoCl_2_ × 6 H_2_O: 0.152 g, CaCl_2_ × 2 H_2_O: 0.1 g, ZnCl_2_ × 4 H_2_O: 0.085 g, CuCl_2_: 0.005 g, AlCl_3_: 0.01 g, H_3_BO_3_: 0.01 g, Na_2_MoO_4_ × 2 H_2_O: 0.01 g, NiCl_2_ × 6 H_2_O: 0.03 g, Na_2_SeO_3_ × 5 H_2_O: 0.0003 g, Na_2_WO_4_ × 2 H_2_O: 0.008 g. Brought final volume to 1L with d H_2_O. Final pH was adjusted to 7 with HCl and NaOH. Vitamin solution (1000X): In 1 L of dH_2_O, dissolved: biotin: 20 mg, folic acid: 20 mg, pyridoxin-HCl: 100 mg, thiamine-HCl × 2H_2_O: 50 mg, riboflavin: 50 mg, nicotinic acid: 50 mg, vitamin B12: 1 mg, D-Ca-panthotenate: 50 mg, p-aminobenzoic acid: 50 mg, lipoic acid: 50 mg. Vitamins were filter sterilized into a sterile anaerobic serum flask (30 mL in 50 mL flask), crimp sealed and degassed by flushing the headspace of the vial for 30 minutes with oxygen free nitrogen at a flow rate of 0.5 LPM through blue cannulas (0.6 mm ID, Microlance, Beckton Dickinson, Franklin Lakes, NJ, USA) equipped with a sterile filter (Minisart, Sartorius, Göttingen, Germany) on the gassing line.

The carbon sources and additions for the monocultures of *Dv, Mb*, and *Mm* as live-control were different. Briefly, 30 mM Na-lactate and 10 mM Na_2_SO_4_ were added for *Dv* monocultures, 100 mM Na-acetate was added for *Mb* monocultures, and 10 mM Na-pyruvate and 682 mM NaCl were added for *Mm* monocultures. *Mm* monoculture headspace was replaced with 2 bar 80%H_2_-20%CO_2_. For the co- and tri-cultures, the carbon source was 30 mM Na-lactate and Na_2_SO_4_ was added at three different levels of 0 mM, 7.5 mM and 15 mM, respectively.

All media were prepared by mixing the basal salt solution and adding 10 mL of the trace element solution and 1 mL Resazurin stock (1g/L) to 1L media. 200 mL of the medium was brought to the boiling point in a 500 mL conical flask and then maintained at 80 °C with a continuous flow of anoxic gas (80% N_2_ + 20% CO_2_) at 0.5 LPM flow rate into the headspace of the flask with a cannula (using a rubber stopper to close off the top opening). After 5 min degassing, vitamin mix stock (0.2 mL into 200mL) and anoxic Cysteine-HCl stock (0.2M, 2 mL into 200mL) were added separately into the medium. The stirring of medium was kept at medium speed with the gas flow as above for 1 hour. The removal of oxygen was verified by a color-shift from pink to colorless by the Resazurin. All gases (BOC, England, UK) used for headspace flushing are run through an oxygen scrubber column (Oxisorb, MG Industries, Bad Soden, Germany), to remove any residual oxygen. All chemicals used are analytic grade or higher (>= 98% purity, Sigma-Aldrich, St. Louis, MO, USA).

The media were prepared in bulk and dispensed into 5 mL per 27 mL Hungate anaerobic culture tubes (Chemglass Life Sciences, Vineland, NJ, USA) in an anaerobic chamber station (MG 500, Don Whitley). This chamber is maintained according to the manufacture’s instruction using N_2_, CO_2_ and H_2_ supplies with an actual gas fraction of 3.14% H_2_ and 5.32% CO_2_, as detected by Micro-Gas Chromatography (GC) (Agilent 490 Micro-GC, Agilent Technologies). The culture tubes had been degassed for 24 hours in the anaerobic chamber. Tubes are closed with degassed blue butyl rubber septa (Chemglass Life Sciences, Vineland, NJ, USA) and crimp sealed. Next, tubes were autoclaved for 15 minutes at 121°C in a desktop autoclave (ST 19 T, Dixon, Wickford, UK). Before inoculation, 50-times concentrated Na_2_S stock solution was added into the medium to achieve a final concentration of 2 mM Na_2_S. All gases used for headspace flushing are run through an oxygen scrubber column (Oxisorb, MG Industries, Bad Soden, Germany), to remove any residual oxygen. All chemicals used are analytic grade or higher (>= 98% purity, Sigma-Aldrich, St. Louis, MO, USA).

### Experimental design and measurements

Co-cultures of *Dv-Mb* and *Dv-Mm* and tri-cultures of *Dv-Mb-Mm* were constructed as shown in supplementary schematic (Fig. S6) and tested for the methane production in three batches of cultivation, each of three weeks duration. In addition, a single round of five weeks’ incubation of co-cultures and tri-culture was also conducted. Individual monocultures were also incubated in the same scheme as alive control. The construction of co- and tri-cultures were done using the inoculum from individual monocultures. *Dv, Mb* and *Mm* were cultivated until late lag phase for 4 days, 21 days and 7days, respectively before inoculation into mixed cultures. The co-cultures and tri-cultures were inoculated with 200 μl individual strain inocula (4% v/v into 5 ml medium). The cultivation was performed in triplicate and incubated at 37 °C for 3 weeks unless there is a specific explanation, and sub-cultured twice. The dilution level for sub-culturing was 5% (v/v).

For co- and tri-culture communities, three treatments of 0 mM, 7.5 mM and 15 mM sulfate were used, with the latter two treatments corresponding to the half and full theoretical amount required to respire 30 mM lactate (see Table 1). Headspace pressure was measured using a needle pressure gauge (ASHCROFT 310, USA) at the beginning and end of each culture batch. At the end of every three weeks, 1.5 ml culture was extracted using 1 mL syringe inside anaerobic chamber and centrifuged at 5500 rpm for 3 min. The biomass and supernatant were separated and stored at −20 °C for further DNA extraction and Ion Chromatography (IC) analyses. Headspace gas fraction was monitored by a Micro-Gas Chromatography (GC), after which culture tubes were opened and the residual culture (~3 ml) was pooled out for pH measurement.

To test the ability of *Dv, Mb* or *Mm* to grow on lactate for methane production in the above setting conditions, individual monocultures of each strain were incubated with medium containing 30 mM Na-lactate as carbon source and 7.5 mM Na-sulfate. In this case, the headspace air fraction for Mm monoculture was the same with the chamber air instead of 80%H_2_-20%CO_2_.

### Optical density, and gas and ion chromatography

Optical density (OD) of the cultures at 600 nm was measured on a daily basis using a spectrophotometer (Spectronic 200E, Thermo Scientific). Gas fraction in tube headspace was detected by Micro-GC with a micro thermal conductivity detector and two columns (Agilent 490, Agilent Technologies). Lactate, acetate, pyruvate and sulfate were measured using Ion Chromatography (Dionex ICS-5000^+^ DP, Thermo Scientific). An analytical anion column with 4 μm ion exchange matrix beads was used according to the following separation conditions. Flow rate: 0.38 ml/min, Pressure: 4300 psi, Column temperature: 30 °C, Eluent: KOH with the gradient in 37 min of 1.5 mM for - 7~0 min pre-run for equilibration, 1.5 mM for 0~8 min (isocratic), increased to 15 mM during 8~18 min, increased to 24 mM during 18~23 min, increased to 60 mM during 23-24 min, and stayed at 60 mM during 24-30 min. IC is equipped with a conductivity-based detector and supplied with MilliQ-water (R > 18.2 Ω) for eluent generation.

### DNA isolation, PCR and quantitative PCR

DNeasy Power Soil Kit (QIAGEN, Germany) was used for isolating genomic DNA according to the manufacturer’s instruction. This genomic DNA isolation kit was formerly sold by MO BIO as PowerSoil DNA Isolation Kit and used for isolating DNA from bacterial-archaeal co-cultures (Ozuolmez *et al.*, 2015). Genomic DNA was quantified using NanoDrop Spectrophotometer (N60, IMPLEN) and stored at −20 for further analyses.

Specific primers were designed for targeting *dsvA* gene of *Desulfovibrio vulgaris* (IMG gene ID: 637121620), *mtaB* gene of *Methanosarcina barkeri* (IMG gene ID: 637699633) and coenzyme F420 hydrogenase of *Methanococcus maripaludis* (IMG gene ID: 2563556008). The specificity of the developed primers was tested and verified by amplifying the DNAs from monocultures of *Dv, Mb* and *Mm* using the Polymerase Chain Reaction (PCR) conditions in the following, respectively. The selected primer pairs to be used in the present study for qPCR detection were Dv_dsvA_1f (5′ -> 3′: TTCGTGTCCGACATCAAGCA) and Dv_dsvA_1R (5′ -> 3′: GTGGGTTTCACCCTCATCGT) for detecting *Dv* (product length: 135 bp), MB_mtaB_f (5′ -> 3′: TGCAAAGAAGACCGGCACTA) and MB_mtaB_r (5′ -> 3′: GAGCAGTCCACCACCAATGA) for detecting *Mb* (product length: 85 bp), and Mm_F420_3F (5′ -> 3′: TCAACAATACACGGCAACGTA) and Mm_F420_3R (5′ -> 3′: GTATCCTTCAGGCGTTCCAA) for detecting *Mm* (product length: 141 bp).

PCR mixtures (in a total volume of 50 μl with ddH2O) contained 1 μl of dNTPs (10 mM; Bio Lab, New England), 4 μl of MgCl_2_ (25 mM; Promega, USA), 2 μl of forward primer (10 μM), 2 μl of reverse primer (10 μM), template DNA (10-20 ng), 10 μl of GoTaq Flexi Buffer (Promega, USA), 2 μl of Bovine serum albumin (4 mg/ml; Bio Lab, New England), 0.25 μl of GoTaq G2 Flexi polymerase (5 U/ml; Promega, USA). PCR mixtures were prepared in bulk volume each time (>500 μl) to minimize the error and the working volume per sample was 25 μl. PCR was conducted using a 96 well thermal cycler (Veriti, Applied Biosystems) with the following procedure: 95 °C for 5 min, 35 cycles of (95 °C for 30 s, an annealing temperature of 60 °C for 30 s, followed by 72 °C for 1 min), and finally 72 °C for 10 min. All PCR products were electrophoresed in 1X TAE buffer on 1.0 % Hi-Res standard agarose gels (AGTC Bioproducts, UK) with 0.01 % GelRed Nucleic Acid Stain (BIOTIUM 10,000X, Hayward CA, USA). DNA band in the Gels was visualized by a gel imaging system (U Genius 3, SYNGENE). Agilent Technologies Stratagene Mx3005P Realtime PCR system and SYBR Green JumpStart Taq ReadyMix were applied (Sigma-Aldrich, USA) for Quantitative PCR (qPCR) analysis. The primers were described in the above.

The total sequence lengths excluding plasmids (bp) were retrieved from NCBI for the use in the present work, which are 3570858 bp (NCBI ID: ASM19575v1), 4533209 bp (NCBI ID: ASM97002v1), and 1746697 bp (NCBI ID: ASM22064v1) for *Dv, Mb* and *Mm*, respectively. Standard DNA template for each strain was diluted using sterile H_2_O (tenfold dilution series) and tested with the unknown samples in one single qPCR run to generate the standard curve. Each standard sample and replicate in the above experimental design were tested in triplicate under qPCR assay with internal reference dye mode (ROX). The correlation coefficients (*R^2^*) of the standard curves were 0.9987 (*Dv*), 0.9973 (*Mb*) and 0.9999 (*Mm*), and the qPCR efficiencies were 96.1% (*Dv*), 96.8% (*Mb*) and 94.4% (*Mm*).

### Mass balance calculations

We perform mass balance calculations based on the assumption that *Methanosarcina barkeri* (*Mb*) and *Desulfovibrio vulgaris* (*Dv*) utilise only the compounded overall reactions 1-2 and 3-5, shown in Table 2, respectively. It is also possible that *Mb* might combine reactions 1 and 2 so to couple acetate reduction with H2 oxidation;

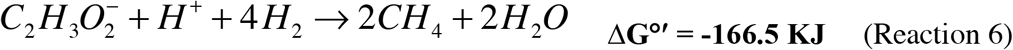

To calculate total methane production in the closed system, we first estimate the amount of acetate and H_2_ available to *Mb*. These compounds can only be produced by *Dv*, through its fermentation pathway, i.e. in reaction 5 from Table 1. We thus calculate produced acetate and H_2_ from observed lactate utilization and the stoichiometry of this reaction. The utilized lactate can be calculated directly from observed lactate at the beginning and end of the cultivation period;

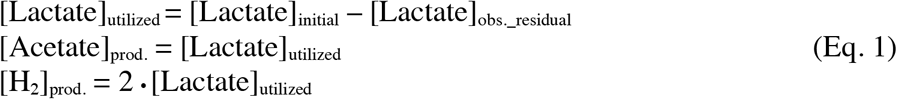

The estimated [Acetate]_prod._ and [H_2_]_prod._ need then be combined with the observed residual levels of these compounds in the system, to estimate the levels that were available to *Mb* ([Acetate]_*Mb*_ and [H_2_]_*Mb*_);

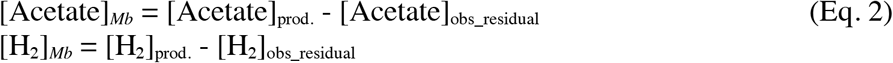

We can now use these values to calculate the estimated stoichiometric, theoretical methane production ([CH_4_]_calc_) by *Mb*, through reactions 1, 2 and 6. The actual amounts of acetate utilized in reactions 2 and 6, as well as the actual amounts of H_2_ utilized in reactions 1 and 6 are unknown. If we assume a full conversion through the three reactions, we would have the following stoichiometric balances;

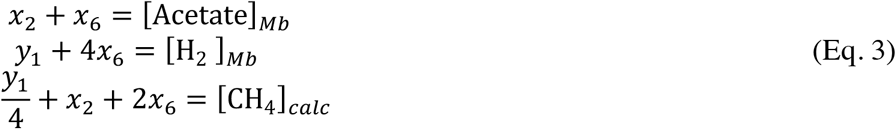

where *x*_i_ and *y*_i_ denote the amounts of acetate and H_2_ utilized in reaction *i*, respectively. These three equalities can then be re-arranged to yield the overall theoretical methane production.

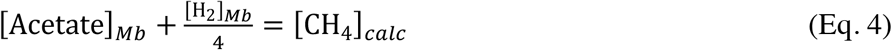

## Author contributions

JC and OSS designed the study and the experiments. JC performed the experiments and analyzed the data. JD, MW, and OSS contributed to analyses of the results and mass balance calculations. All authors contributed to the writing of the manuscript and have given approval to the final version.

